# *Bigh3* is essential for pulmonary fibrosis

**DOI:** 10.1101/2025.09.02.673852

**Authors:** Anthony Altieri, Yashar Aghazadeh Habashi, Yein Chung, Erika E. McCartney, Ronen Schuster, Mark Abovsky, Faizah S. Abdullah, Shaheed Hakim, Sam Cooper, Igor Jurisica, Boris Hinz, Matthew B. Buechler

## Abstract

Transforming growth factor beta (TGF-<u>β)</u>-induced <u>g</u>ene-<u>h</u>uman, clone <u>3</u> (BIGH3) has been implicated as a biomarker of lung fibrosis. However, it is unknown if BIGH3 plays a functional role in fibrosis pathogenesis. To address this question, we used *in silico*, *in vitro* and *in vivo* approaches. We found that *BIGH3*/*Bigh3* is upregulated in human lung fibrosis and mouse models of pulmonary fibrosis. We next generated a novel *Bigh3* knockout (*Bigh3*^−/−^) mouse and found that while these animals exhibited lung architecture and immune cellularity that is broadly equivalent to wild-type mice, they were protected from lung fibrosis in response to bleomycin administration. *In silico* modeling suggested that BIGH3 can bind to integrin alpha v (ITGAV). *In vitro* co-culture systems revealed that activated human lung fibroblasts can elicit *BIGH3* expression from human monocyte-derived macrophages. Last, macrophages elicited collagen expression from lung fibroblasts in a manner that is *Bigh3*-dependent. Collectively, these data suggest that *Bigh3* is a product of fibroblast-macrophage interactions that is essential for the pathogenesis of lung fibrosis, possibly via interactions with ITGAV.

## Introduction

Fibrosis, the pathological scarring of organs, accounts for more than one third of disease-related deaths worldwide (Zeisberg and Kalluri, 2013). Idiopathic pulmonary fibrosis (IPF) is characterized by progressive lung scarring that reduces lung compliance and gas exchange, resulting in compromised breathing and, eventually, death (Murray, 2016; Rockey et al., 2015). IPF is incurable and the two currently approved therapeutics (pirfenidone and nintedanib) are not universally effective and lack defined mechanisms of action (Somogyi et al., 2019; Cottin et al., 2018). Pulmonary fibrosis is driven by activated fibroblasts – called myofibroblasts – that generate overabundant extracellular matrix (ECM) proteins, including collagens, to produce the stiff scar leading to lung dysfunction (Somogyi et al., 2019; Tsukui et al., 2024). Drivers of fibroblast activation are manifold but include cytokines secreted by macrophages (Mϕ) (Aran et al., 2019), most notably the transforming growth factor beta (TGF-β) family of cytokines (Yue et al., 2010). Clinical trials targeting the TGF-β pathway have failed to cure fibrosis in part because TGF-β exerts pleiotropic effects aside from its role in fibrosis in tissue development and in controlling functional aspects of cells within the immune system (Chapman et al., 2020; Nishimura, 2009).

An in-depth characterization of the cellular dynamics in mouse models of fibrosis has the potential to yield new therapeutic targets for treatment of pulmonary fibrosis in humans. We have previously proposed that focusing on genes that tune TGF-β signaling may represent novel treatment pathways for fibrosis (Altieri et al., 2024). For example, TGF-β Inducible (*TGFBI*), referred to here as TGF-<u>β</u>-induced <u>g</u>ene-<u>h</u>uman, clone <u>3</u> (BIGH3) (LeBaron et al., 1995), is produced downstream of TGF-β signaling in humans and mice (SKONIER et al., 1992; Altieri et al., 2024). BIGH3 contains a signal peptide, enabling its secretion, four fasciclin 1 domains known to mediate binding to ECM proteins, and an arginine-glycine-aspartic acid (RGD) motif, which serves as a recognition site for integrins (Kawamoto et al., 1998; SKONIER et al., 1992, 1994). Periostin (POSTN), another fibrosis-associated matrisomal and ECM protein, is a paralog of BIGH3 that shares high sequence homology but lacks that RGD domain required for integrin recognition (Mosher et al., 2015; Hynes and Naba, 2012). BIGH3 is upregulated in the lungs of patients with lung fibrosis and has been proposed as a biomarker for IPF (Xu et al., 2021), yet its functional role in pulmonary fibrosis has not been definitively elucidated. In this study, we used genetic tools, *in silico* and *in vitro* approaches to show that BIGH3 operates as a key driver of pulmonary fibrosis, possibly by binding to integrin alpha v (ITGAV), a key integrin in mediating fibrosis (Henderson et al., 2013), and is a product of fibroblast-Mϕ crosstalk.

## Materials and methods

### Mice and bleomycin administration

Mice were bred and maintained in groups of 1-5 animals per cage at The Centre for Phenogenomics (TCP; Toronto, ON, CAN). Males and females from 6-12 weeks old were used and were age/sex matched in experiments. Tgfbi_del_wt, Tgfbi_del_het or *Dpt*^IresCreERT2^;*Cthrc1*^FLOX^ (that did not receive tamoxifen) were used as wild type / control animals and were bred and maintained at TCP. Tgfbi_del_hom mice were used as *Bigh3*^−/−^ (See below). *Col1a1-*GFP mice were a generous gift from David Brenner at UC San Diego (Yata et al., 2003).

Bleomycin was administered via passive inhalation as in (Song et al., 2025; Cai et al., 2024). Briefly, animals received 50 microliters of saline or Bleomycin (Fresenius Kabi) at 0.25 units per kilogram (U/kg) three times on Monday – Wednesday – Friday (one dose per day) such that the total dose was 0.75 U/kg. To do so, mice were deeply anesthetized with isoflurane in an induction chamber. The front teeth of the mice were then hanged on a horizontal wire apparatus (ETI-MSE-01; Kent Scientific; CT, US) that ensured an open mouth and enabled nose-cone anesthesia during inhalation. The tongue was gently pulled out with a cotton swab. A soft pipette tip was inserted into the airway and used to deliver the bleomycin preparation or sterile saline (control). When the pipette tip created a seal in the airway, the prepared solutions were inhaled into the lung. Once the dosing solution was inhaled out of the pipette tip, the mice were removed from the wire and placed on a heat pad and monitored continuously until fully conscious and ambulatory. All experiments were performed under TCP and University of Toronto approved protocols.

### Generation of BIGH3 knockout mouse

The mouse line C57BL/6J-Tgfbi<em1Tcp> (referred to here as *Bigh3^−/−^*mice) was generated at TCP using methods previously described (Gertsenstein and Nutter, 2021). Briefly, these mice were generated by introducing a 779-bp deletion that affected exons 7 and 8. This deletion corresponded to optimal loxP sites for the generation of cell-specific *Bigh3* knockout mice. Exon 7 is a critical exon in the *Tgfbi* gene (ENSMUST00000045173). This allele was generated by electroporating Cas9 ribonucleoprotein complexes with single guide RNAs having spacer sequences of CATAAGGACTCTGCTAACCG targeting the 5’ side and CAAGAGCTTGGACGAATCAG targeting the 3’ side of a critical region in ENSMUSE00000245131. This resulted in a 779-bp deletion of Chr13:56775342 to 56776120 (GRCm39). The repair templates were not incorporated during repair.

### Analysis of publicly-available single cell RNAseq data

Human single cell sequencing data and metadata were downloaded from Gene Expression Omnibus (GEO) with accession number GSE136831 (Adams et al., 2019), using metadata containing additional cell information such as patient diagnosis (control, IPF, or COPD). Subsequent data analysis was performed in Rstudio (v.4.4.1) using Seurat (v.5.3.0). Only control and IPF patients were included in our analysis (COPD samples were excluded). Ensembl gene IDs were mapped to Hugo Gene Nomenclature Committee (HGNC) symbols using the R package BioMart (v.2.60.1). Multiplets (denoted in the metadata) and low-quality cells expressing less than 1000 unique genes, with more than 20% being mitochondrial genes, were removed. Gene expression of the remaining 136,891 cells was log-normalized, and the top 2,000 highly variable genes were detected with the *FindVariableFeatures*(). Data were scaled with mitochondrial gene percentage regressed out, and principal component analysis was used for dimensionality reduction. Nearest neighbors were found, and cells were clustered using the first 30 principal components and a resolution of 0.5, respectively. Uniform manifold approximation and projection (UMAP) was used for two-dimensional data visualization. Cell type annotations were provided in the metadata, with marker validation shown as in (Adams et al., 2019).

Mouse single cell sequencing data were downloaded from the GEO with accession number GSE141259 (Strunz et al., 2020) and were analyzed in Rstudio (v.4.5.0) using Seurat (5.3.0). The raw data set had previously undergone quality control filtering, resulting in a final data set with 29,297 cells provided by the authors. Gene expression of the remaining cells was log-normalized, and the top 3,000 highly variable genes were detected with the *FindVariableFeatures()* function. Data were scaled and principal component analysis was used for dimensionality reduction. Batch effects were corrected by integration with the R package Harmony (v.1.2.3). Nearest neighbors were found, and cells were clustered using the Harmony-corrected dimensions and a resolution of 0.8, respectively. UMAP was used for two-dimensional data visualization. Cell type annotations were denoted in the supplementary metadata, with marker validation shown in (Strunz et al., 2020).

Differential abundance (DA) analysis was performed using the R package MiloR (v.2.5.1), by which significant neighborhoods were identified using spatial FDR correction. Data were visualized with Beeswarm plots and overlaying DA results onto previously generated UMAPs.

### Whole lung RNA isolation, cDNA synthesis and quantitative PCR

Left lung lobe was added to the QIAGEN (ON, CAN) Power Bead tubes (Ceramic 2.8 mm, Cat. 13114-50) containing 350 uL of RLT Plus lysis buffer (QIAGEN) and homogenized using bead mill homogenizer (OMNI International; OK, USA) with the following program: 3000 rpm, 3 cycles, for 50 sec/cycle. Total RNA was extracted from the homogenized tissue using RNeasy Plus Mini Kit (QIAGEN, Cat. 74136) in accordance with manufacturer’s guideline and quantified by Nanodrop Eight Spectrophotometer (Thermofisher; MA, USA). Reverse transcription was performed with 200 ng of RNA ABI advanced kit included RNase inhibitor (ABI Cat. 4374966). qPCR was performed on cDNA product diluted with RNase-free water (1:100) using Taqman Gene Expression Assay probe for the genes: *Col1a1* (Mm00801666_g1) and *Tgfbi* (Mm01337605_m1). *Gapdh* (Mm99999915_g1) was used as the reference gene. Taqman Fast Advanced Master Mix (Thermofisher, 4444963) was used to dilute each probe (1:10) and performed with the following cycling parameters: 50°C for 2 min, 95°C for 20 sec, and then repeating 40 cycles, 95°C for 1 sec, 60°C for 20 sec. All qPCR reactions were conducted in duplicate in a MicroAmp optical 384-well reaction plate using QuantStudio 6Pro (Applied Biosystems; MA, USA). The average of threshold cycle (Ct) of each target gene was normalized to *Gapdh*.

### Suspension Cytometry by Time of Flight (CyTOF)

Mice were sedated using aerosolized isoflurane. Animals were injected retro-orbitally using an insulin syringe (BD; NJ, USA; Cat. 305932) with 3ug CD45.2 147Sm (Standard Biotools; ON, CAN; Cat. 3147003B) in 100-125uL of sterile saline. After three minutes under sedation, mice were euthanized by cervical dislocation and lungs were excised. Lungs were then weighed, minced and digested using the digestion mix as in (Buechler et al., 2021). Specifically, tissues were digested in 3mL of digestion mix for 8 minutes in a 37C water bath, then supernatant was removed with a pipette, filtered through a 70uM cell strainer (Falcon; NY, USA; Cat. 352350) into cold PBS with 2% FBS and 0.5M EDTA. Next, 3 mL of digest mix was added to remaining tissue and incubated for 12 minutes and supernatant was removed as before. Tissue was then incubated in 3mL of digest mix for 15 minutes and supernatant was removed as before. Red blood cells were lysed with Ack lysis buffer (Gibco; MA, USA; Cat. A10492-01) and cells were counted using a ViCell XR (Beckman Coulter; CA, USA) or using a hemocytometer.

3 x 10^6^ cells were aliquoted into 4 mL polystyrene V-bottom tubes (Diamed; ON, CAN; catalog # STK 8550) for CyTOF staining and were washed with CyTOF staining medium (CSM; PBS containing 1% BSA). Cells were Fc blocked at room temperature (RT), followed by incubation with metal-conjugated antibodies for 30 minutes at RT. The cells were then washed once with CSM. Subsequently, the cells were washed with PBS alone and incubated with 200 μL of 1 μM cisplatin solution (BioVision; CA, USA; catalog #1550–1000) for five minutes at RT for viability staining. Cisplatin was quenched by adding CSM (containing 1% BSA). Cellular DNA was labelled by incubating cells with 1 mL of 100 nM of iridium (Fluidigm; CA, USA; Catalog #201192B) in PBS containing 0.3% saponin and 1.6% formaldehyde for 1h at RT. The stained cells were washed, resuspended in 100 μL fetal calf serum containing 10% DMSO, and stored at –80 °C until acquisition.

Pellets of stained cells were transferred on ice to SickKids-UHN Flow and Mass cytometry Facility for acquisition. Cells were resuspended in Maxpar Cell Acquisition Solution containing EQ beads (diluted 1:10) and filtered through cell strainer cap tubes (Falcon; Catalog# 352235).

Cells were acquired on a third-generation Helios mass cytometer (Fluidigm) at 100-250 events per second. The acquired files are normalized with the preloaded normalizer algorithm on CyTOF software version 6.7.

### CyTOF analytical pipeline

CD45– and CD45.2-populations were concatenated by group (i.e. *Bigh3*^+/+^ and *Bigh3*^−/−^ mice). These populations were then normalized to the lowest number of cells in each group and concatenated to generate a representative cell population. UMAPs were generated using the UMAP_R plugin for FlowJo 10.9.0.

### Histology and pathological scoring

Formalin fixed whole lung or lung lobes were processed in a Tissue Tek VIP 6 tissue processor (Sakura Finetek; CA, USA), paraffin-embedded, 4 µm-thick sections were collected onto charged slides (Assure, Epic Scientific; NJ, USA) and stained with hematoxylin & eosin or Masson’s trichrome in a Tissue Tek Prisma auto-stainer (Sakura Finetek). Slides were scanned at 20X using an Olympus VS120 slide scanner equipped with a Hamamatsu ORCA-R2 C10600 digital camera (Evident, Japan).

All slides were randomly numbered in a blinded fashion and scored by an anatomical pathologist, using the modified Ashcroft scale (Hbner et al., 2008). Briefly, the lungs were inspected microscopically with a 20x objective, following a raster-like pattern throughout the sections. Foci with predominant trachea or bronchial mucosa were excluded. Each 20x focus was graded, summarized, and then divided by the total number of fields to obtain a fibrotic index for the lung.

### Bulk RNAseq and analysis

Whole lung RNA was isolated as described above. Quality of total RNA was isolated was assessed using an RNA nano chip on an Agilent Bioanalyzer (Agilent; CA, USA). For each sample, 200 ng RNA was used to prepare sequencing libraries using trueseq stranded total RNA library preparation kits with RiboZero as per the manufacturer’s instructions (Illumina; CA, USA). Pooled samples volumetrically pooled and sequenced on an Illumina Next Seq 550 sequencer for 150 paired end read-cycles. Only protein-coding genes were retained for downstream analysis using a custom list of mouse protein-coding genes generated from NCBI (**Supplemental Table 1**). Differential gene expression analysis was performed using the DESeq2 package (v.1.49.3). A regularized log (rlog) transformation was applied to normalized counts, and sample-level variability was assessed by principal component analysis (PCA). Gene set enrichment analysis (GSEA) was conducted using the clusterProfiler (v.4.17.0) and fgsea (v.1.35.6) R packages. Normalized enrichment scores from GSEA were visualized in custom plots using ggplot2 (v.3.5.2).

### Culture of mouse bone marrow Mϕ

Mouse Mϕ. Tibia and femur pairs were harvested from *Bigh3*^−/−^ and *Bigh3*^+/+^ mice, decapped to expose the bone marrow, placed open side down in a PCR tube perforated by an 18.5G needle, and centrifuged at 12,000 RPM for 20 seconds. Bone marrow cells were resuspended in cold PBS-EDTA. Red blood cells were lysed by Ack Lysing Buffer (Gibco; cat#A10492-01) for 3 minutes on ice. After, cells were transferred to a 15 mL conical tube and Ack buffer was neutralized with 5 mL of RPMI containing 10% FBS (Gibco; cat#A31607-01), 1% Penicillin/Streptomycin (Gibco; Cat#15140122), and 1% L-Glutamine (Gibco; Cat#25030081), hereafter referred to as R10. Cells were counted and 1×106 cells were plated in R10 containing 10 ng/mL recombinant murine M-CSF (Peprotech; NJ, USA; Cat# 315-02) for 2 days. After 2 days, cells were washed twice with PBS (Gibco; Cat#14190144) and cultured in R10 containing 20 ng/mL CSF1 and 10 ng/mL IL-4 (Cat # 214-14) and 10 ng/mL IL-13 (Peprotech; Cat 210-13) (M_2-IL4+IL13_) for 3 days.

### Co-culture of human activated fibroblasts and Mϕ and single cell RNAseq analysis

Human peripheral blood mononuclear cells (PBMCs; STEMcell Technologies; BC, CAN) were enriched for CD14⁺ monocytes using a magnetic separation kit (Miltenyi Biotec, Germany), according to the manufacturer’s instructions. Isolated monocytes were cultured in macrophage base medium (RPMI-1640 supplemented with 10% fetal bovine serum, 1% penicillin–streptomycin, and 2 mM l-glutamine; all from Gibco) in the presence of recombinant human Mϕ colony stimulating factor (M-CSF, 50 ng ml⁻¹; PeproTech) for 6 d to generate unpolarized Mϕ. Polarization was induced by culturing unpolarized Mϕ for an additional 48 h with either lipopolysaccharide (LPS, 100 ng ml⁻¹; Sigma–Aldrich; MO, USA) and interferon-γ (IFN-γ, 20 ng ml⁻¹; PeproTech) to generate M_1-IFNG_ Mϕ, or interleukin-4 (IL-4, 20 ng ml⁻¹; PeproTech) to generate M_2-IL-4_ Mϕ.

Human activated fibroblasts were cultured from the CCD-8Lu lung fibroblast cell line (ATCC) at passage 3 on tissue culture-treated plastic dishes, maintained in DMEM supplemented with 10% fetal bovine serum and 1% penicillin–streptomycin and combined with fibroblasts treated as above with recombinant human TGF-β1 (2 ng ml⁻¹; PeproTech) for 48 h before co-culture. Mϕ were co-cultured with activated fibroblasts at a 1:5 Mϕ-to-fibroblast ratio for 72 h in Mϕ base medium.

All single-cell data were combined into a single Scanpy object for analysis. Low quality cells were filtered out based on two criteria: fewer than 6,000 detected genes and mitochondrial gene content exceeding 10%. Gene expression values were normalized to 10,000 counts per cell and log-transformed. The top 2,000 highly variable genes were identified and scaled. Principal component analysis (PCA) was then performed on the scaled variable genes. The top 40 principal components were used for neighborhood graph construction and graph-based clustering (resolution = 0.2), as well as UMAP dimensionality reduction. All preprocessing and analysis steps were conducted using functions from the Scanpy package (e.g. pp.filter_cells, pp.normalize_total, pp.highly_variable_genes, pp.scale, tl.pca, pp.neighbors, tl.umap, tl.leiden) with default parameters unless otherwise specified. Mϕ were defined as cells enriched for MRC1-high cells (MRC1 > 5 counts per cell), fibroblasts were identified as cells expressing COL1A1-high (COL1A1 > 9 counts per cell). Differential gene expression analysis was carried out using the Wilcoxon rank-sum test, comparing Mϕ monoculture to co-cultures with fibroblasts in either direct contact or transwell culture.

### Co-culture of fibroblasts and Mϕ for live cell imaging

Mouse M_2-IL4+IL13_ Mϕ were isolated and polarized as described above. Supernatant from M2 Mϕ was collected after centrifugation at 1600 rpm. Mouse lung fibroblasts were digested as previously described. 10e6 whole lung cells were plated on a 10mm dish in D10 for 24 h then washed 2x with sterile PBS. Cells were cultured in D10 for 7 days under 80% confluency. Cells were then washed 2x with sterile PBS, then trypsinized with 0.25% Trypsin-EDTA (1X; Gibco 25200-056). Cells were incubated with Fc Block (1:1000; Biolegend; cat#101302) for 5 minutes on ice, washed with PBS, then incubated with CD45 microbeads (Miltenyi Biotec; Cat#130-052-301) and CD31 microbeads (Miltenyi Biotec; Cat#130-097-418) for 15 minutes on ice before magnetic-activated cell sorting (MACS)-based cell separation to deplete CD45+ and CD31+ cells. Fibroblast identify was verified by flow cytometry. 1×10^4^ lung fibroblasts/well were added to a 96-well plate and 200 uL supernatant from *Bigh3*^−/−^ and *Bigh3*^+/+^ M_2-IL4+IL13_ Mϕ supernatant was added to each well. The 96-well plate was placed on the Incucyte SX5 (Sartorius; NY, USA), a Live-Cell Imaging and Analysis Instrument. *Col1a1*-GFP+ integrated intensity was measured over 7 days by imaging *Col1a1*-GFP+ fibroblasts using the phase and green fluorescence channels at 10X magnification in an incubator at 37°C with 5% CO2.

### Protein-protein interaction analysis

TGFBI or POSTN were used to query IID database v.2025-05 (https://iid.ophid.utoronto.ca/iid) (Kotlyar et al., 2021). Gene Ontology Biological Process was used to annotate individual proteins, as per legend. Protein interaction networks were visualized in NaViGaTOR v.3.0.19 (Brown et al., 2009; Pastrello et al., 2014) and the final figure was combined with a legend in Adobe Illustrator v.29.7.1 using the SVG file. Their interaction was modeled using the docking software MEGADOCK version 4.1.1

(https://www.bi.cs.titech.ac.jp/megadock/ppi.html (Hayashi et al., 2018). The visualization and interpretation of the docking results were carried out using ChimeraX version 1 10.1 (2025-07-24) visualization software ((https://www.rbvi.ucsf.edu/chimerax (Meng et al., 2023).

### Data and code availability

Publicly-available scRNAseq data were obtained from the Gene Expression Omnibus (GEO) under accession numbers GSE136831 (human) and GSE14125 (mouse). All code used for analysis and figure generation is available at https://github.com/BuechlerLab/BIGH3-LUNG-FIBROSIS-2025.

## Results and Discussion

### BIGH3/Bigh3 is upregulated in human IPF and mouse models of lung fibrosis in fibroblasts and Mϕ

To explore if BIGH3 is associated with lung fibrosis, we utilized publicly available single-cell RNA-sequencing (scRNA-seq) data from patients with IPF (Adams et al., 2020) and a mouse model of bleomycin-induced pulmonary fibrosis (Strunz et al., 2020). Uniform manifold approximation and projection (UMAP) of scRNA-seq data from human lungs revealed that *BIGH3* is expressed by both fibroblasts and Mϕ (**Figure 1a and Supplemental Figure 1a**). During IPF, a phenotypic shift occurred in Mϕ clusters, with an increase of interstitial Mϕ, as identified by *SPP1* and *CD36* (Leach et al., 2020; Adams et al., 2020), and the development of a myofibroblast cluster identified by expression of *COL3A1*, *CTHRC1*, *COL7A1,* and *TNC* (Buechler et al., 2021) (**Figure 1b and Supplemental Figure 1b**) consistent with other reports (Aran et al., 2019). These clusters, which emerged with the development of fibrosis, displayed high expression of *BIGH3* (**Figure 1c**).

**Figure 1.**
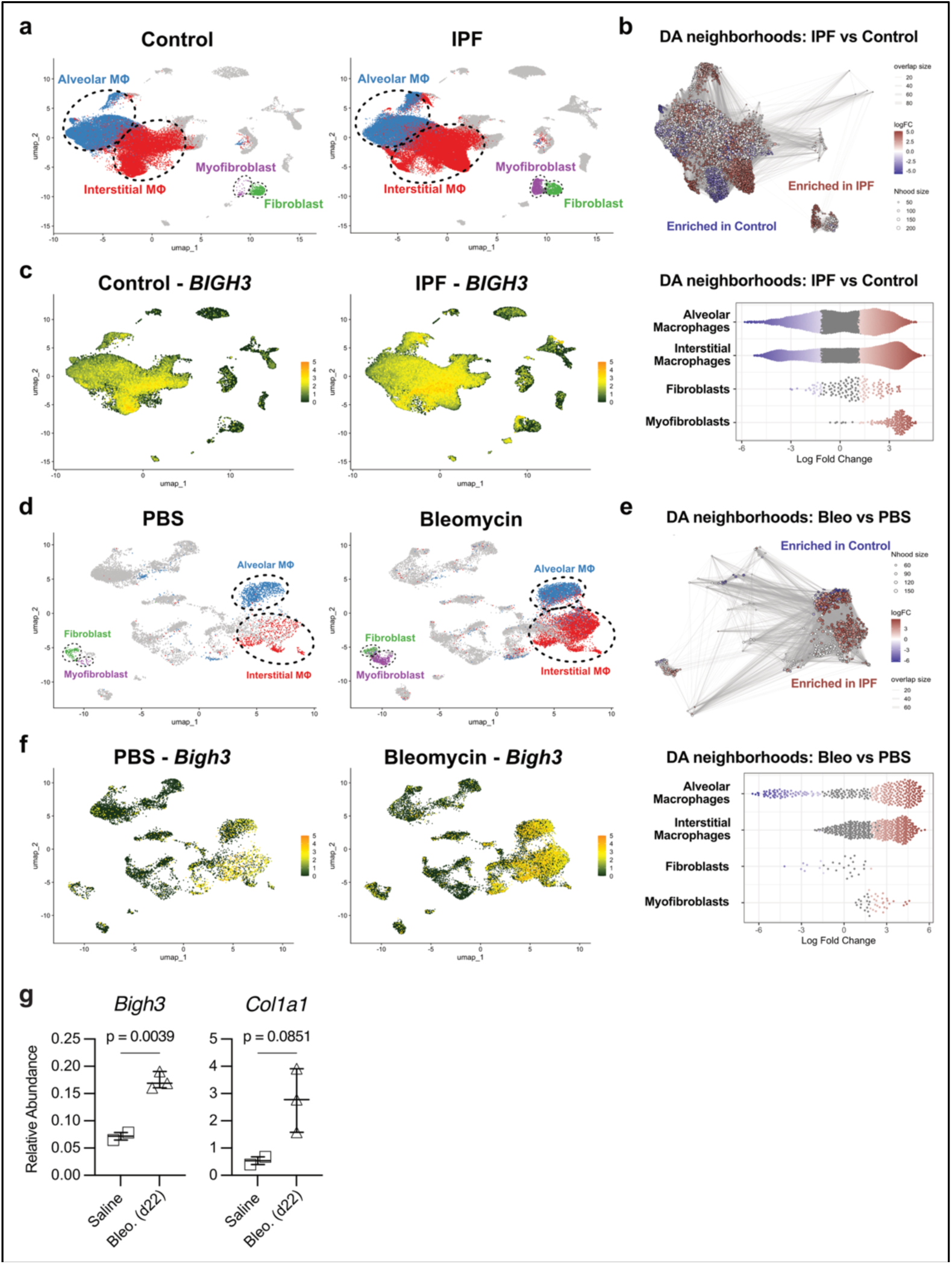
*BIGH3/Bigh3* is upregulated in human IPF and mouse models of pulmonary fibrosis. **a**, UMAP of human lung scRNA-seq data from control and IPF patients, highlighting alveolar MΦ (blue), interstitial MΦ (red), fibroblast (green), and myofibroblast (purple) cell populations. **b,** Milo differential abundance analyses of Mϕ and fibroblast neighborhoods across control and IPF conditions, with a graph highlighting enriched regions (top) and Beeswarm plot showing log fold changes by cell type (bottom). **c,** *BIGH3* expression across UMAP embeddings in control and IPF lungs. **d,** UMAP plots of scRNA-seq data from lungs of mice receiving PBS or bleomycin PITI, highlighting alveolar MΦ (blue), interstitial MΦ (red), fibroblast (green), and myofibroblast (purple) cell populations. **e,** Milo differential abundance analyses of Mϕ and fibroblast neighborhoods across PBS and bleomycin conditions, with a graph highlighting enriched regions (top) and Beeswarm plot showing log fold changes by cell type (bottom). **f,** *Bigh3* expression across UMAP embeddings in PBS– and bleomycin-administered mouse lungs. **g,** abundance of *Col1a1* and *Bigh3* in whole lung 24 d after bleomycin. Representative of 3 or more biologically independent experiments. Symbol represents one mouse with bar at mean and plots showing minimum and maximum (whiskers) and median (centre line). *p<0.05 as determined by Student’s t-test.

In mice, pulmonary administration of bleomycin elicits a defined inflammatory response after 2–7 days (d), characterized by immune cell infiltration into the lung, followed by an extended fibrotic phase after 14–24 d (Tsukui et al., 2020; Peyser et al., 2019; Selman and Pardo, 2001; Tsukui et al., 2024; Cai et al., 2024; Song et al., 2025). In a mouse model of bleomycin-induced pulmonary fibrosis, fibroblasts, myofibroblasts, and interstitial Mϕ expressed *Bigh3* (**Figure 1d-f and Supplemental Figure 2**). In these datasets, *POSTN*/*Postn* was restricted to myofibroblasts and vascular endothelial cells and not expressed in Mϕ (**Supplemental Figures 1e & 2e**). These analyses reveal that *BIGH3/Bigh3*-expressing myofibroblasts and interstitial Mϕ arise in the lung during IPF and in mouse models of pulmonary fibrosis.

To validate the results from scRNA-seq data analysis, we exposed mice to three doses of bleomycin via passive inhalation (Song et al., 2025; Cai et al., 2024). At 21 d, we observed evidence of fibrosis, including increased expression of *Collagen 1a1 (Col1a1)* in whole-lung lysates and a concomitant significant increase in *Bigh3* expression compared to mice that received saline as a control (**Figure 1g**). These data indicate that fibroblasts and Mϕ express *BIGH3/Bigh3* in humans and mice, and that BIGH3/*Bigh3* becomes more abundant during pulmonary fibrosis.

### Bigh3 deficient mice exhibit normal lung structure and immune composition in the steady state

We next set out to elucidate a functional role of BIGH3 in pulmonary fibrosis using a constitutive genetic knockout (KO) mouse we generated. These mice, referred to hereafter as *Bigh3^−/−^* mice, harbored a 779-base pair deletion that affected exons 7 and 8 of *Bigh3* (**Figure 2a**). The lungs of *Bigh3^−/−^* mice did not express *Bigh3* under steady-state conditions, indicating the fidelity of the model (**Figure 2b**). Consistent with previous reports, adult *Bigh3^−/−^* mice exhibited no obvious gross defects in lung architecture but were smaller than their littermate controls by whole-body weight (Ahlfeld et al., 2016) (**Figure 2c-d**). In the steady state, there was no difference in the absolute number or number per milligram of cells per lung between *Bigh3^−/−^* and *Bigh3^+/+^* control animals (**Figure 2e-f**).

**Figure 2.**
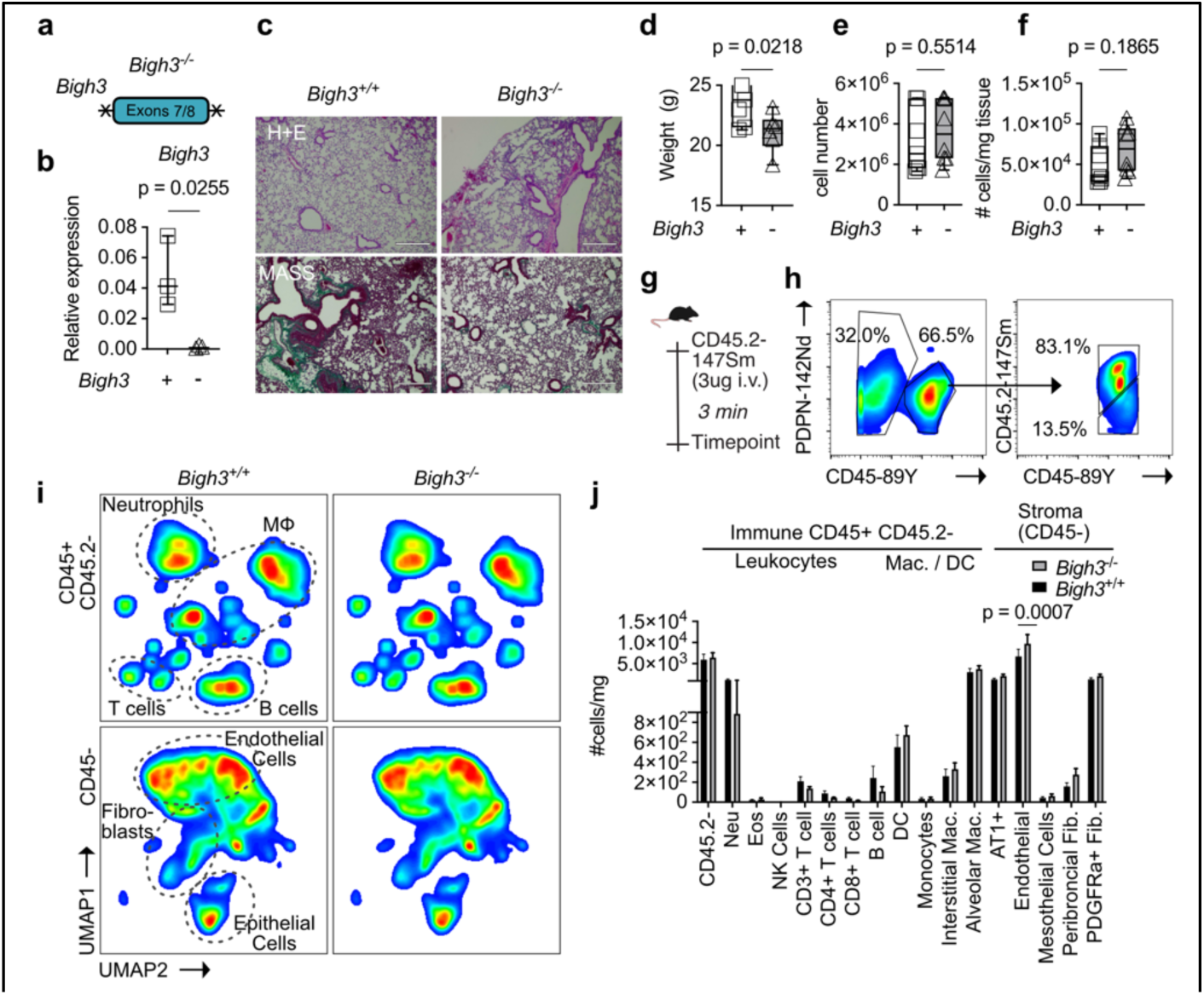
*Bigh3*^−/−^ mouse exhibits normal lung structure and immune composition in the steady state. **a**, schematic of *Bigh3*^−/−^ transgenic mouse, **b,** abundance of *Bigh3* in the lungs of *Bigh3*^+/+^ and *Bigh3*^−/−^ mice, **c,** representative images of *Bigh3*^+/+^ and *Bigh3*^−/−^ and hematoxylin and eosin (top) and Masson’s Trichome stained (bottom) stained lungs, **d,** whole body weight, **e,** number of cells and **f,** number of cells per milligram of tissue in the lungs of *Bigh3*^+/+^ and *Bigh3*^−/−^ mice, **g,** graphical schematic of retro-orbital CD45.2-147Sm antibody injection, **h,** representative gating of CD45-, CD45+, CD45.2-, and CD45.2+ cells, **i,** UMAPs of CD45+CD45.2-(top) leukocytes and CD45-stromal cells in the lungs *Bigh3*^+/+^ and *Bigh3*^−/−^ mice **j,** enumeration of leukocyte and stromal cells in the lungs of *Bigh3*^+/+^ and *Bigh3*^−/−^ mice, data shown as mean +SEM. All comparisons between *Bigh3*^−/−^ and *Bigh3*^+/+^ mice were determined by a student’s t-test (d-f) or multiple unpaired t test (j; only endothelial cells displayed a significant difference between groups). Each symbol represents 1 mouse in b, d-f. Data are from 1 (b) or 2 (d-j) experiments with greater >= 7 mice per group (d-j).

To determine the cellular composition of the lung in *Bigh3^−/−^*mice, we employed cytometry by time-of-flight (CyTOF) analysis, which included 25 parameters selected to identify immune and stromal cell populations in the lung (**Supplemental Tables 2 and 3**). To label lung-resident immune cells, an antibody specific for CD45.2 conjugated to a 147Sm isotope was intravenously injected and animals were euthanized 3 minutes later (**Figure 2g**). This approach revealed that in the lung, on average, 85-90% of cells in *Bigh3^−/−^*and *Bigh3^+/+^* controls are CD45.2-labeled and therefore in the bloodstream or exposed to blood flow, consistent with previous literature (Barletta et al., 2012) (**Figure 2h**).

To determine if *Bigh3*^−/−^ mice exhibited differences in immune cell and fibroblast populations compared to control animals, multi-dimensional CyTOF data were projected in UMAP space. Analysis within these UMAPs revealed no difference in immune or stromal populations between *Bigh3^−/−^* and *Bigh3^+/+^* mice (**Figure 2i and Supplemental Figure 3**). Individual immune cell populations including B cells, T cells, Nk1.1+ cells, monocytes, alveolar Mϕ, interstitial Mϕ, dendritic cells, eosinophils, and neutrophils were tabulated, and no difference was observed in the proportion of live CD45+CD45.2-cells or the absolute number per milligram of lung tissue between groups. No difference in the percentage of live cells or the number per milligram of tissue between fibroblast subtypes (Tsukui et al., 2020; Buechler et al., 2021), mesothelial (Buechler et al., 2019), or epithelial cells was observed. *Bigh3^−/−^* mice did exhibit a statistically significant increase in CD31+ endothelial cells relative to *Bigh3^+/+^* controls (**Figure 2j-k**). These data indicate that the structure and cellular composition of the lung in adult mice is broadly similar between *Bigh3^−/−^* mice and *Bigh3^+/+^* animals under steady-state conditions.

### Bigh3 is essential to develop fibrosis after bleomycin administration

BIGH3 has been suggested to be essential for the expression of *Collagen 1a1* (*COL1A1)* downstream of TGF-β signaling in fibroblasts (Yang et al., 2022). However, the mode(s) of BIGH3 action under these conditions require additional mechanistic studies and *in vivo* validation. As we revealed that *BIGH3*/*Bigh3* is upregulated in the context of pulmonary fibrosis and that *Bigh3*^−/−^ mice exhibit no overt lung defects in the steady state, we examined whether deletion of *Bigh3* affected lung fibrosis in mice.

To test the role of *Bigh3* in pulmonary fibrosis, we administered bleomycin to *Bigh3^−/−^* and *Bigh3^+/+^* control mice (Song et al., 2025; Cai et al., 2024). *Bigh3^−/−^* maintained body weight comparable to saline treated controls. In contrast, *Bigh3^+/+^* animals lost significant body weight after bleomycin treatment compared to animals that received saline, as well as compared to *Bigh3^+/+^* mice that were administered bleomycin (**Figure 3a**), indicating that *Bigh3* may influence responses to inhaled bleomycin across the inflammatory and fibrotic phases of bleomycin treatment. After 21 d, the lungs of *Bigh3^−/−^* mice exhibited significantly less *Bigh3* and *Col1a1* RNA compared to *Bigh3^+/+^*animals (**Figure 3b,c**). Histologically, *Bigh3^−/−^* mice that received bleomycin harbored less collagen deposition near airways and blood vessels relative to *Bigh3^+/+^* animals, as determined by Masson’s Trichrome staining. Consistent with these observations, *Bigh3^−/−^*mice displayed a significantly lower Modified Ashcroft score as a measurement of lung fibrosis, as determined by blinded scoring by a pathologist, compared to *Bigh3^+/+^* mice (**Figure 3d-e**). These data indicate that *Bigh3* can function as a regulator of the fibrotic response in an experimental model of lung fibrosis.

**Figure 3.**
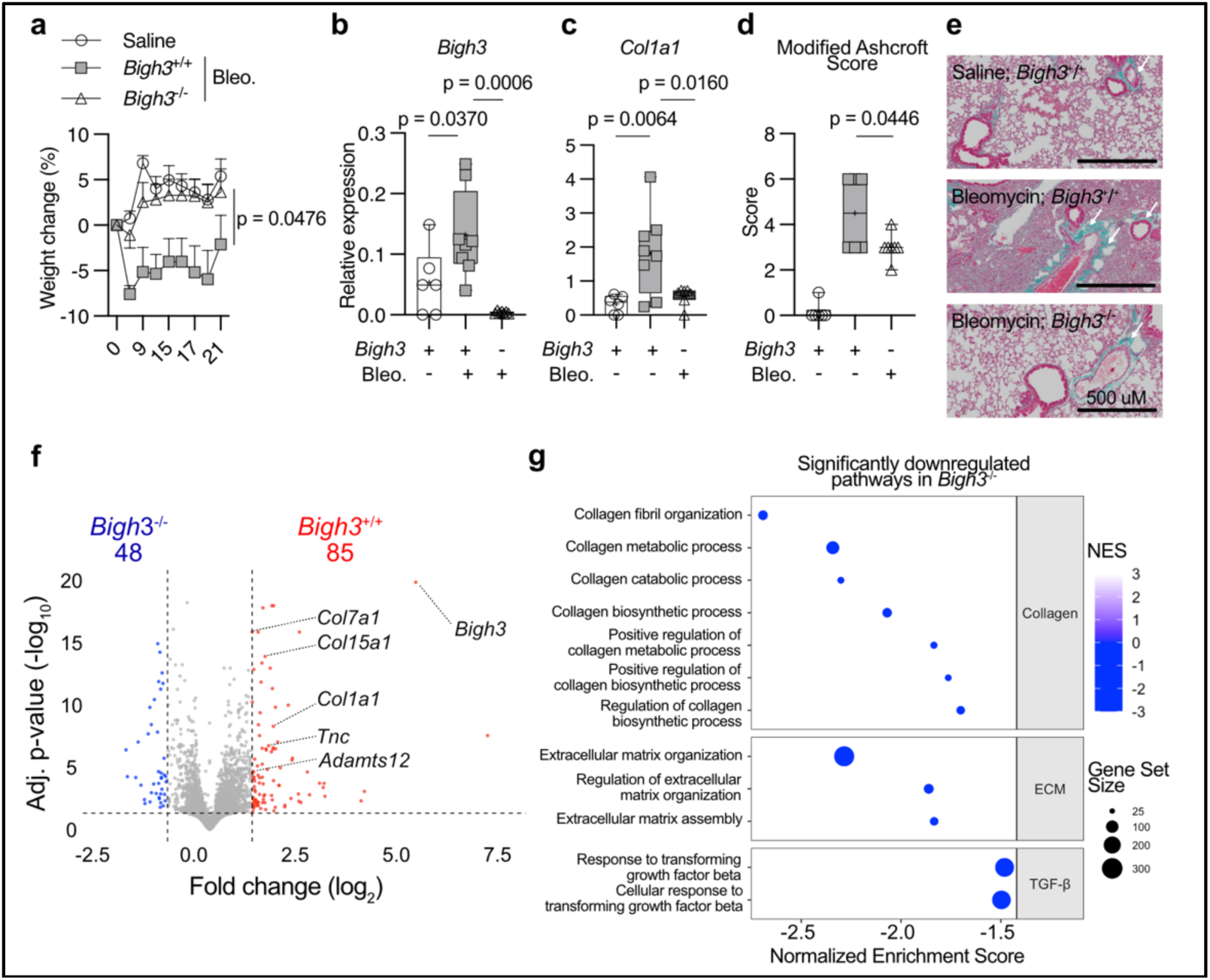
*Bigh3* is required for fibrosis to develop after bleomycin administration. **a**, % weight change in saline-or bleomycin-administered *Bigh3*^−/−^ and *Bigh3*^+/+^ mice **b,** *Bigh3* and **c,** *Col1a1* abundance in saline-or bleomycin-administered *Bigh3*^−/−^ and *Bigh3*^+/+^ mice**, d,** modified Ashcroft scores in saline-or bleomycin-administered *Bigh3*^−/−^ and *Bigh3*^+/+^ mice **e,** representative images of saline-or bleomycin-administered *Bigh3*^−/−^ and *Bigh3*^+/+^ mice **f,** volcano plot of differentially expressed genes in saline-or bleomycin-administered *Bigh3*^−/−^ and *Bigh3*^+/+^ mice **g,** GSEA bubble plot with pathways downregulated in the lungs of *Bigh3*^−/−^ compared to Bigh3^+/+^ mice. All comparisons between *Bigh3*^−/−^ and *Bigh3*^+/+^ mice were determined by a student’s t-test or multiple unpaired t test (a-d). Each symbol represents 1 mouse and data are shown as mean +SEM in b-d. Data are from 3 (a) or 2 (b-e) independent experiments with greater >= 6 mice per group or 1 experiment with n = 3 mice per group (f-g).

To better understand the dynamics of lung fibrosis in the absence of *Bigh3* after 21 d, we performed bulk RNA-seq on whole lungs from mice treated with saline and *Bigh3^+/+^* and *Bigh3^−/−^* mice that received bleomycin (**Supplemental Figure 4a)**. Genes associated with fibrotic lungs, such as collagens and other ECM components, significantly increased in *Bigh3^+/+^* mice that received bleomycin compared to *Bigh3^+/+^* mice that received saline (**Supplemental Figure 4b and Supplemental Table 4**). Consistent with differential gene expression, geneset enrichment analysis (GSEA) showed increased pathways associated with ECM and collagen deposition in *Bigh3*^+/+^ mice that receive bleomycin compared to those that were administered saline (**Supplemental Figure 4c**).

To evaluate the effect BIGH3 has on lung fibrosis, we next compared *Bigh3^+/+^* and *Bigh3^−/−^* mice that received bleomycin. Compared to *Bigh3^−/−^* mice, *Bigh3^+/+^* mice expressed significantly increased levels of genes associated with fibrosis, including *Col1a1*, *Col7a1, Col15a1, Tnc, and Adamts12* (**Figure 3f and Supplemental Table 5**). GSEA analysis further indicated that pathways associated with collagen organization, metabolism, and synthesis, as well as ECM deposition and TGF-β-mediated signaling pathways were diminished in the lungs of *Bigh3*^−/−^ mice compared to *Bigh3*^+/+^ controls (**Figure 3g**). In sum, these data demonstrate that after bleomycin administration, mice lacking *Bigh3^−/−^* exhibited diminished fibrosis and reduced gene expression in the lungs associated with decreased TGF-β signaling and ECM deposition compared to control animals that received bleomycin.

### BIGH3 can bind to ITGAV, is a product of fibroblast-Mϕ interactions and is required for maximal Mϕ-induced collagen expression from fibroblasts

Previous studies have suggested that integrin signaling, specifically ITGAV, is essential for pulmonary fibrosis pathogenesis (Henderson et al., 2013). BIGH3 has been shown to bind to αv integrin (Peng et al., 2022). We validated this observation *in silico* by examining predicted protein-protein interaction (PPI) partners of BIGH3 using the integrated interactions database (IID) (Kotlyar et al., 2021). This approach revealed that *BIGH3* is predicted to interact with more than 20 other proteins, including several collagens and integrins, such as ITGAV (**Figure 4a and Supplemental Table 5**). POSTN was predicted to interact with several collagen family proteins, like BIGH3, but did not exhibit associations with integrins (**Supplemental Figure 5 and Supplemental Table 6**). Consistent with associations between integrins in the literature and our PPI analysis, integrin-associated signaling pathways were decreased in the lungs of bleomycin-administered *Bigh3*^−/−^ mice compared to *Bigh3*^+/+^ controls (**Figure 4b**). We conducted molecular docking and computational modeling to provide additional detail for the interaction between ITGAV and BIGH3. The molecular docking analysis revealed several contact points connecting exons 29 and 30 of ITGAV (amino acids 991-1045) to exons 7 through 12 of BIGH3 (amino acids 288-533) (**Figure 4c and Supplemental Table 7 and Supplemental Video 1**).

**Figure 4.**
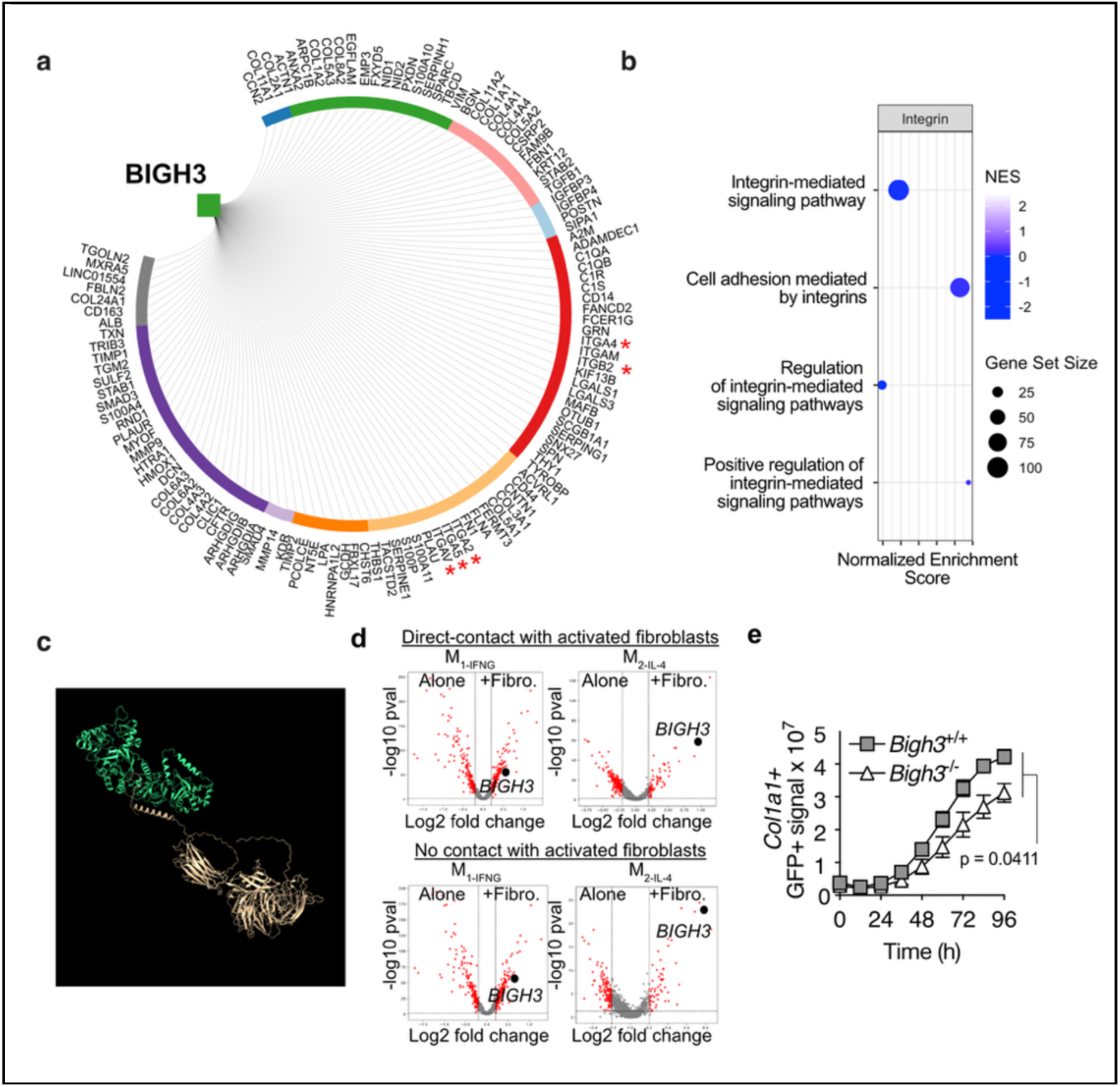
Fibroblast-Mϕ interactions can elicit BIGH3-dependent ECM deposition from fibroblasts. **a**, Protein-protein interaction (PPI) diagram for BIGH3. Node color indicates GO Biological Process Node: Cell Aggregation (Blue), Cellular Component Organization of Biogenesis (Green), Developmental Process (Salmon), Immune System Process (Red), Locomotion (Pale marigold), Metabolic Process (Orange), Rhythmic Process (Lavendar), Signaling (Purple), Growth (Light blue), Uncategorized (Gray). Integrins denoted by red asterisk. **b,** GSEA bubble plot demonstrating significantly downregulated integrin-associated pathways in the lungs of *Bigh3*^−/−^ compared to *Bigh3*^+/+^ controls **c,** Highest-ranking model of ITGAV-BIGH3 biding, where the gold corresponds to ITGAV (the receptor), and the green corresponds to BIGH3 (the ligand). **d,** fibroblast-Mϕ coculture Mϕ bulk RNA-seq **e,** *Bigh3*^+/+^ and *Bigh3*^−/−^ M_2-IL-4/IL-13_ coculture with *Col1a1*-GFP+ lung fibroblasts. All comparisons between *Bigh3*^−/−^ and *Bigh3*^+/+^ mice were determined by a student’s t-test (d). Each symbol represents the average +/-SEM from 3 technical replicates from one experiment, representative of two experiments. p<0.05 as determined by students t test.

Our analysis of scRNA-seq mouse and human pulmonary fibrosis datasets indicated that fibroblasts and Mϕ were the predominant source of *BIGH3/Bigh3*. To dissect whether *Bigh3* can be a product of fibroblast-Mϕ crosstalk, we setup heterocellular co-cultures. Herein, human monocyte-derived Mϕ from peripheral blood were polarized to ‘M_1-IFNG_’, or ‘M_2-IL-4_’ phenotypes, then co-cultured directly or separated by a transwell insert with human lung myofibroblasts and subsequently recovered for scRNA-seq analysis (**Supplemental Figure 6a**). These data revealed that co-culture of human Mϕ with activated fibroblasts shifted Mϕ towards a ‘M_2-IL-4_’ phenotype, independent of initial polarization state (**Supplemental Figure 6b-c**). *BIGH3* was significantly upregulated by both Mϕ activation states upon co-culture with activated fibroblasts in a manner that was independent of cell contact (**Figure 4d and Supplemental Table 8**). These data reveal that activated fibroblasts can elicit *BIGH3* from Mϕ.

We next sought to determine if Mϕ-derived BIGH3 induced functional changes in fibroblasts that could influence fibrosis pathology. Our data demonstrated that *Col1a1* was significantly suppressed in the lungs of *Bigh3*^−/−^ mice compared to *Bigh3*^+/+^ *in vivo* (**Figure 3c, f**), suggesting that *Col1a1* serves as a surrogate metric of BIGH3 activity. To this end, fibroblasts derived from the lungs of *Col1a1-*green fluorescent protein (GFP) mice (Yata et al., 2003) were cultured in medium conditioned by M_2-IL-4/IL-13_ polarized Mϕ derived from the bone marrow of *Bigh3^−/−^* or *Bigh3^+/+^* animals. These cultures were imaged 2 times daily for 96 h to quantify GFP expression (*Col1a1 promoter activity*). This approach revealed significantly more GFP in cultures containing *Bigh3^+/+^*M_2-IL-4/IL-13_ supernatant compared to cultures with *Bigh3^−/−^*M_2-IL-4/IL-13_ conditioned medium (**Figure 4e**). These data support a model in which Mϕ can elicit the expression of *Col1a1* from fibroblasts in a manner that is significantly influenced by *Bigh3*.

We employed genetic tools and *in silico* and *in vitro* approaches to reveal that *Bigh3* is indispensable for lung fibrosis and is a product of fibroblast-Mϕ crosstalk. Our findings support that *Bigh3* production by Mϕ augments ECM deposition by ITGAV-expressing myofibroblasts. Cell-type-specific transgenic tools that enable selective abrogation of *Bigh3* are required to characterize the source(s) of *Bigh3 in vivo* that influence fibrosis pathogenesis and pinpoint whether this protein influences ECM deposition, inflammation or both stages of lung fibrosis. Additional studies are also needed to understand whether *Bigh3* impacts the cellular composition of the fibrotic lung, acts to alter the function of fibrotic cells in this space, or operates via combination of these actions. *Postn* knockout mice are also protected from developing lung fibrosis in response to bleomycin (Naik et al., 2012; Uchida et al., 2012), though our protein-protein analysis suggests *Bigh3* and *Postn* may operate through distinct mechanisms of action. Therapeutic approaches targeting matrisomal proteins such as *Bigh3* and *Postn* (Hynes and Naba, 2012), and both may yield effective therapies for fibrotic indications, including, but not limited to, pulmonary fibrosis. These may work via a two-factor authentication method as previously proposed (Altieri et al., 2024) or via new, conceptually novel mode(s) of action that will be elucidated by future studies.

## SUPPLEMENTAL FIGURES

**Supplemental Figure 1.**
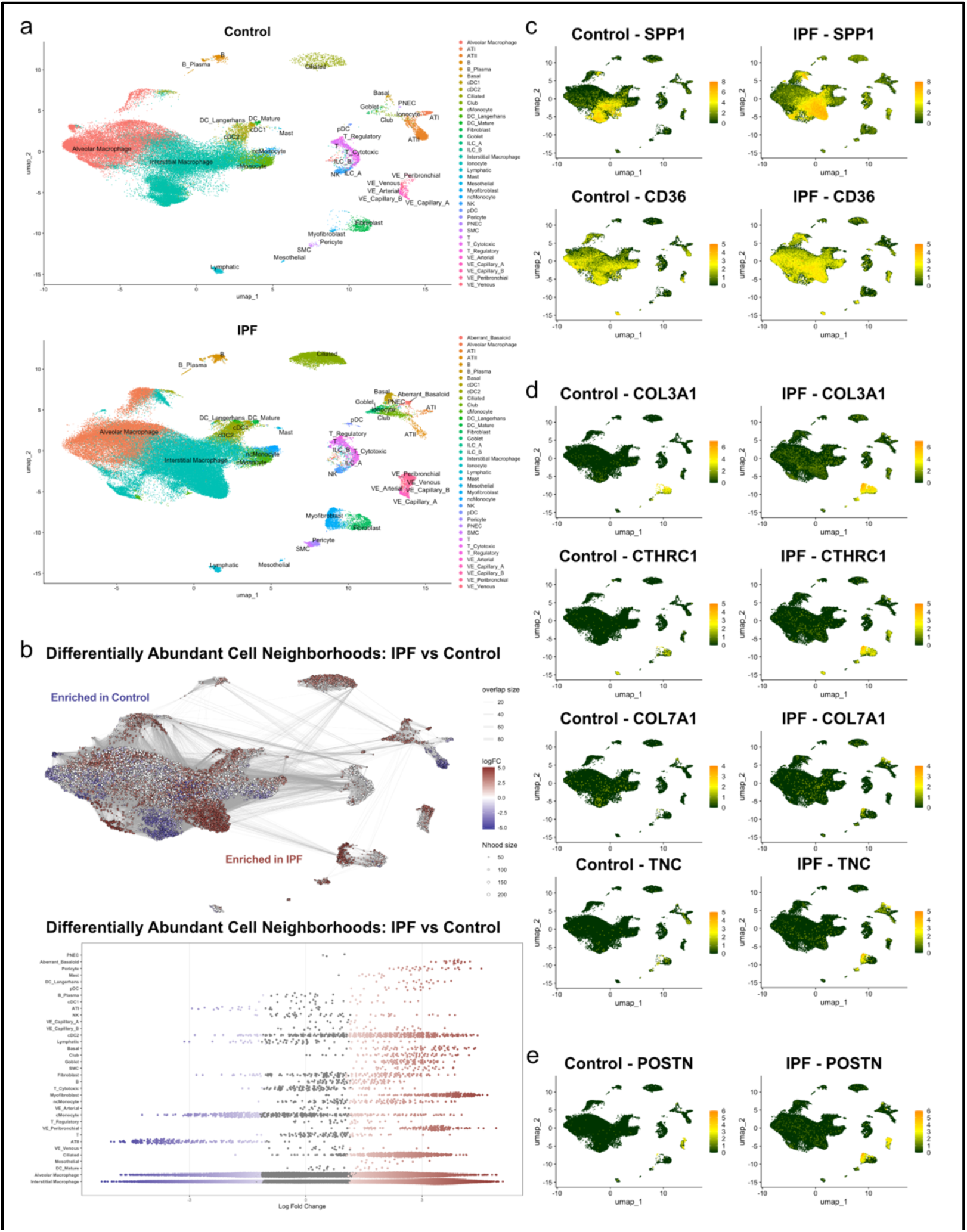
*BIGH3* and *POSTN* are upregulated in human IPF and mouse models of pulmonary fibrosis. **a**, UMAP of human lung scRNA-seq data from control (top) and IPF (bottom) patients, with annotated cell clusters. **b,** Milo differential abundance analyses of all cell neighborhoods across control and IPF conditions, with a graph highlighting enriched regions (top) and Beeswarm plot showing log fold changes by cell type (bottom). **c,** *SPP1* and *CD36* expression across UMAP embeddings in control and IPF lungs. **d,** *COL3A1*, *CTHRC1*, *COL7A1*, and *TNC* expression across UMAP embeddings in control and IPF lungs. **e,** *POSTN* expression across UMAP embeddings in control and IPF lungs.

**Supplemental Figure 2.**
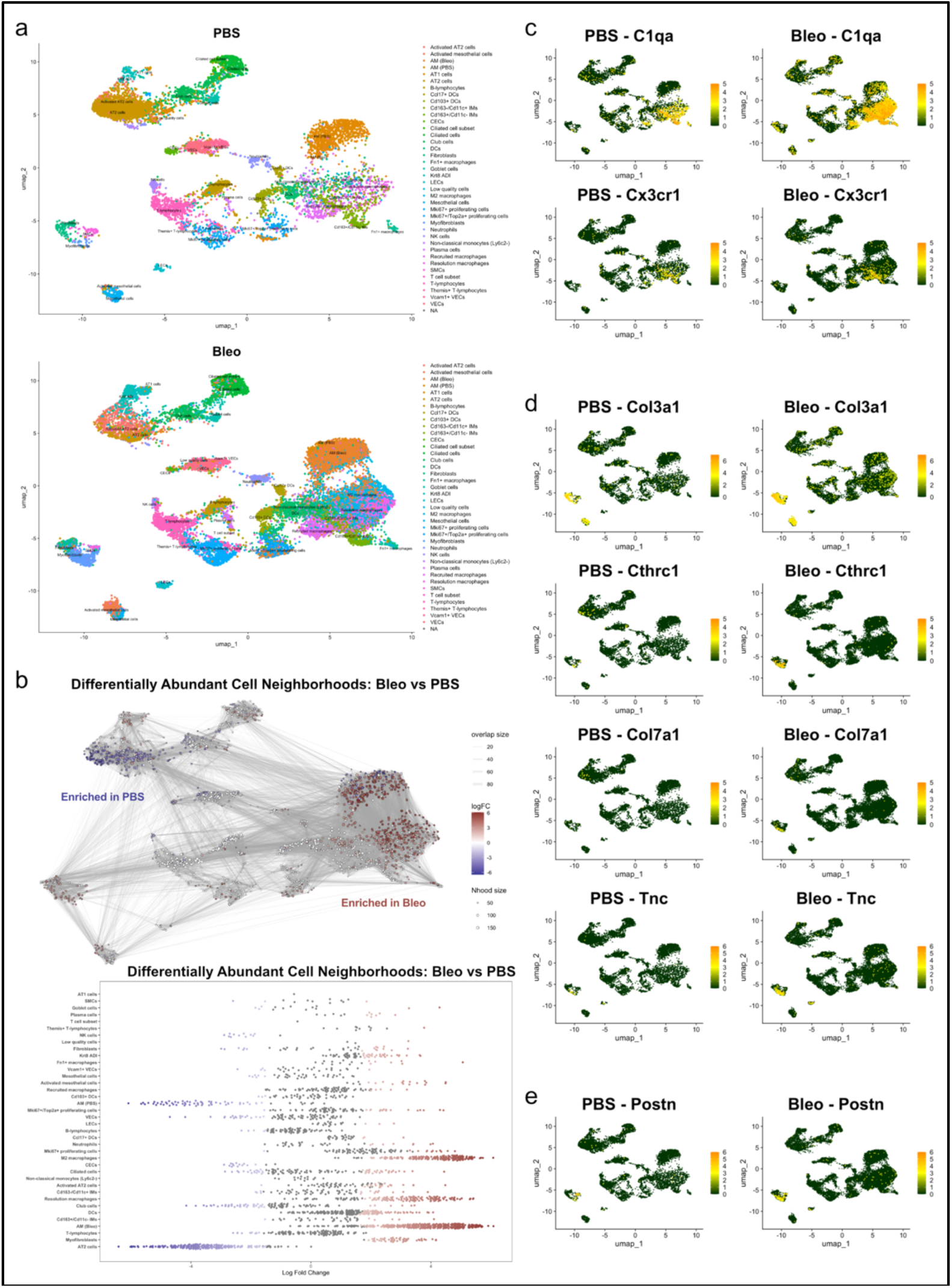
*Bigh3* and *Postn* are upregulated mouse model of pulmonary fibrosis. **a**, UMAP plots of scRNA-seq data from lungs of mice receiving PBS (top) or bleomycin (bottom). **b,** Milo differential abundance analyses of all cell neighborhoods across PBS and bleomycin conditions, with a graph highlighting enriched regions (top) and Beeswarm plot showing log fold changes by cell type (bottom). **c,** *C1qa* and *Cx3cr1* expression across UMAP embeddings in control and IPF lungs. **d,** *Col3a1*, *Cthrc1*, *Col7a1*, and *Tnc* expression across UMAP embeddings in control and IPF lungs. **e,** *Postn* expression across UMAP embeddings in control and IPF lungs.

**Supplemental Figure 3.**
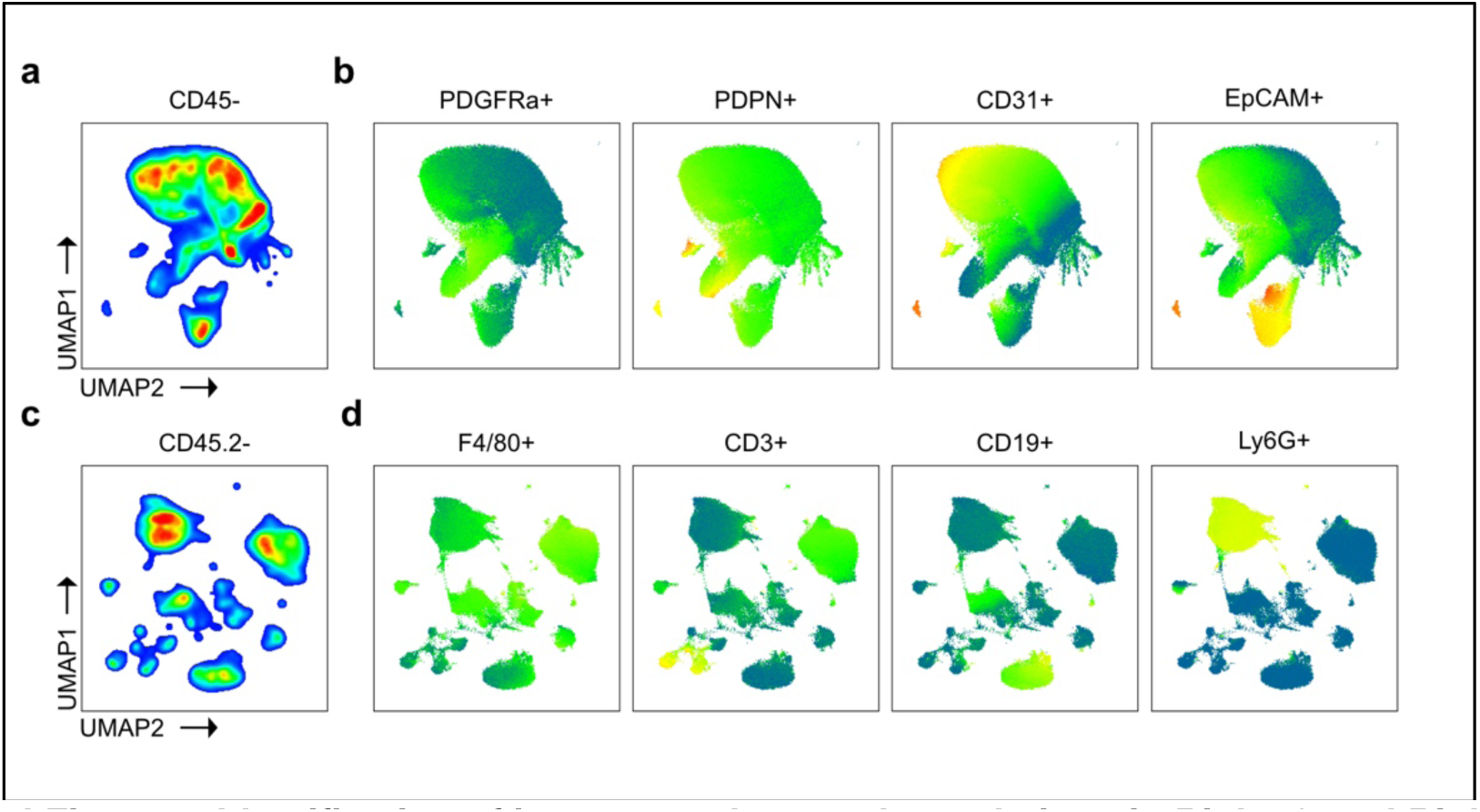
Identification of immune and stromal populations in Bigh3^+/+^ and Bigh3^−/−^ mice. **a**, UMAP of CD45-stromal cells in the lungs of *Bigh3*^+/+^ and *Bigh3*^−/−^ mice. **b**, heatmap statistic plots of PDGFRa+, PDPN+, CD31+, EpCAM+ of CD45-stromal cells in the lungs of *Bigh3*^+/+^ and *Bigh3*^−/−^ mice. **c**, UMAP of CD45.2-leukocytes in the lungs of *Bigh3*^+/+^ and *Bigh3*^−/−^ mice. **d**, heatmap statistic plots of F4/80+, CD3+, CD19+, and Ly6c+ leukocytes cells in the lungs of *Bigh3*^+/+^ and *Bigh3*^−/−^ mice.

**Supplemental Figure 4.**
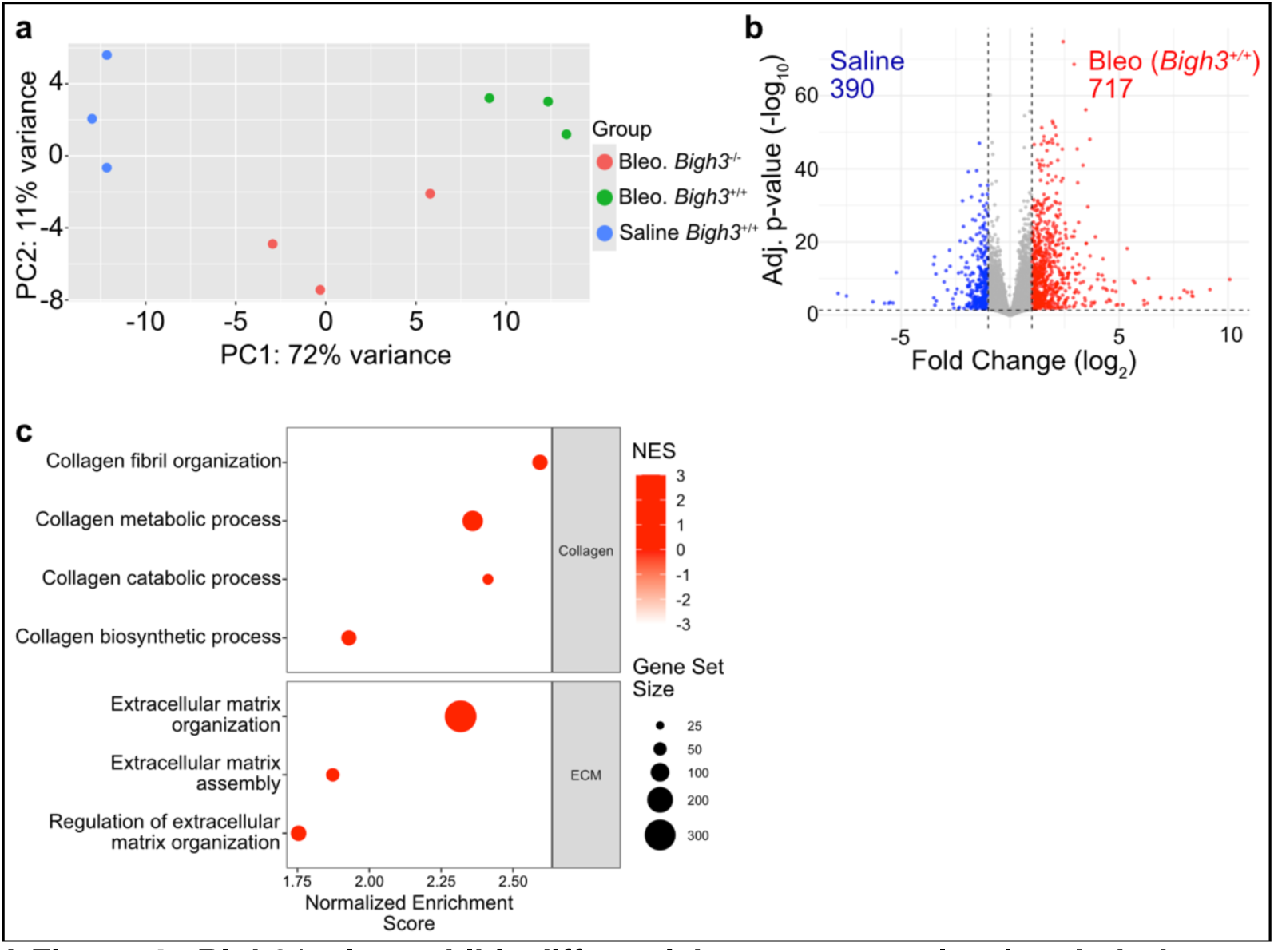
*Bigh3^−/−^* mice exhibit differential gene expression in whole lungs compared to *Bigh3^+/+^* mice that receive bleomycin. **a**, principal component analysis of bulk RNA-seq profiles showing distinct transcriptional segregation of Saline *Bigh3*^+/+^, Bleo. *Bigh3*^+/+^, and Bleo. *Bigh3*^−/−^ groups. **b,** volcano plot of differentially expressed genes in lungs of saline– and bleomycin-administered *Bigh3*^+/+^ mice. **c,** GSEA bubble plot demonstrating significantly upregulated collagen– and ECM-associated pathways in lungs of *Bigh3*^+/+^ mice compared to *Bigh3*^−/−^ mice.

**Supplemental Figure 5.**
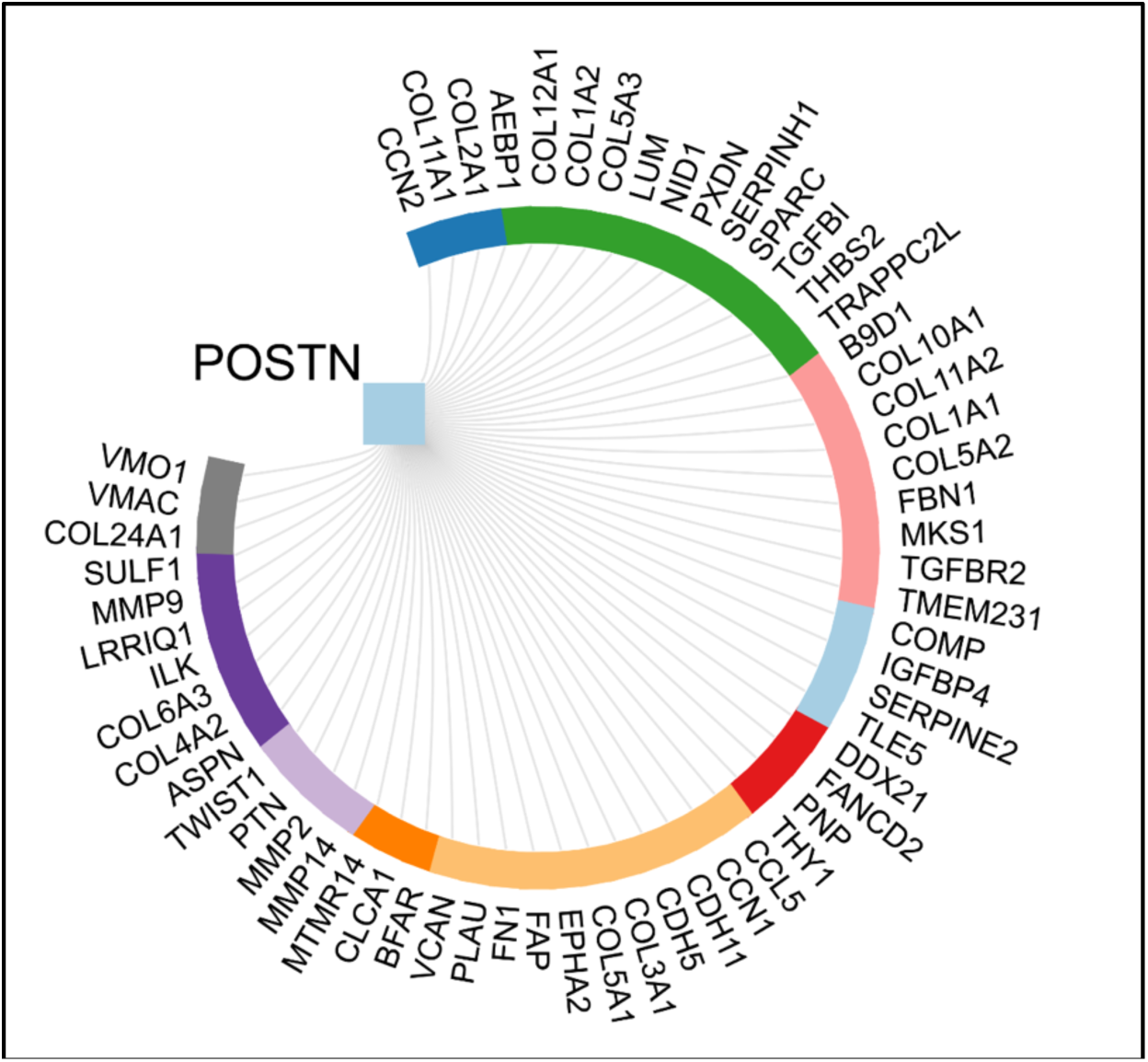
POSTN is not predicted to interact with integrins. Protein-protein interaction (PPI) diagram for POSTN. Color indicates GO Biological Process Node: Cell Aggregation (Blue), Cellular Component Organization of Biogenesis (Green), Developmental Process (Salmon), Immune System Process (Red), Locomotion (Pale marigold), Metabolic Process (Orange), Rhythmic Process (Lavendar), Signaling (Purple), Growth (Light blue), Uncategorized (Gray).

**Supplemental Figure 6.**
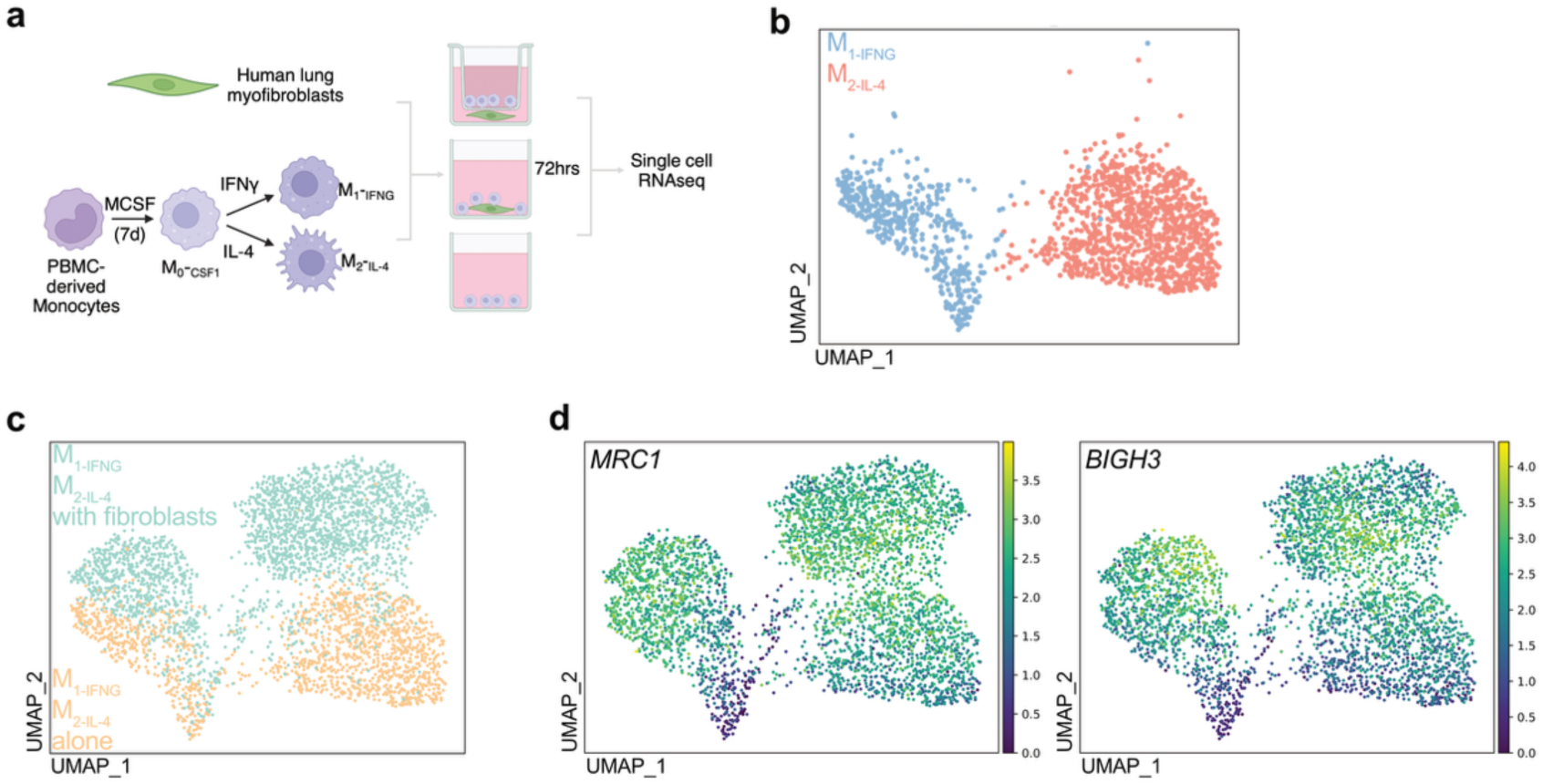
Fibroblast-Mϕ crosstalk elicits BIGH3 expression by Mϕ. **a**, Experimental schematic. **b,** Mϕ from Mϕ monocultures after polarization with cytokines (top right). **c,** Mϕ (green) from Mϕ from fibroblast co-cultures (orange). **d,** Featureplots showing *MRC1* and *BIGH3* by Mϕ in Mϕ monocultures and from fibroblast co-cultures.

## Acknowledgements

We thank the Buechler Lab for constructively reading and editing this manuscript. We are grateful to the staff at the Centre for Phenogenomics for animal husbandry and colony maintenance assistance. We thank the Centre for Immune Analytics in the Department of Immunology at the University of Toronto for assistance with the Incucyte SX5 platform. We would also like to thank Dr. Mohit Kapoor and the Schroeder Arthritis Institute for assistance with bulk RNA-seq. CyTOF was performed at the SickKids’ Centre for Advanced Single Cell Analysis and supported by the SickKids Foundation and a Canadian Foundation for Innovation Fund John Evans Fund Leaders’ Infrastructure Grant to Cynthia J Guidos (#37562). CJG’s work was supported by a Canadian Institutes of Health Research Project Grant (FRN 165973).

## Author Contributions

Conceptualization: BH, MBB

Investigation / Methodology: AA, YAH, YC, EEM, FSA, MA, RS, SH, SC, IJ, MBB

Visualisation: AA, YAH, YC, EM, MA, SC, IJ, MBB

Funding acquisition: IJ, BH, MBB

Project administration: IJ, BH, MBB

Supervision: SC, IJ, BH, MBB

Writing – original draft: AA, YAH, MBB

Writing – review & editing: AA, YAH, EEM, YC, IJ, SC, BH, MBB

